# Novel ciliary protein TRIM8 is a multifunctional workhorse during mitosis

**DOI:** 10.1101/2025.03.28.646005

**Authors:** Utsa Bhaduri, Eleonora Di Venere, Gabriella Maria Squeo, Giorgia Gemma, Francesco Tamiro, Rosario Avolio, Emanuela Senatore, Lucia Salvemini, Rossella Di Paola, Danilo Licastro, Ilaria Iacobucci, Paolo Salerno, Antonio Feliciello, Maria Monti, Vincenzo Giambra, Giuseppe Merla

## Abstract

TRIM8 is an E3 ubiquitin ligase that functions as both a tumour suppressor and an oncoprotein. Earlier, we reported that TRIM8 interacts with key regulators of mitotic spindle assembly, and that TRIM8 knockdown results in mitotic delay and aneuploidy. In this study, we implemented a multi-omics strategy with differential transcriptomic (single-cell RNA sequencing or scRNA-seq), translatomic (polysome profiling with RNA-seq), and proteomic (LC-MS/MS) approaches to elucidate the involvement of TRIM8 in different levels (transcription, translation, post-translation) and stages (G0/G1, S, G2/M) of mitotic cell cycle regulation and progression. With the aid of differential transcriptomic (scRNA-seq) and proteomic (LC-MS/MS) approaches, we show that depletion of *TRIM8* perturbs the canonical “Cell Cycle Control of Chromosomal Replication” pathway and demonstrate that TRIM8 negatively regulates the expression of *TOP2A*, known to be essential for genomic integrity. We also show that TRIM8 downregulation induces substantial alterations in the translation activity of cells and results in the upregulation of polysome-bound *MALAT1* lncRNA by means of significant changes in polysome profiling coupled with RNA-sequencing. Moreover, we unveil endogenous TRIM8 as a novel ciliary protein that co-localizes with CEP170, required for ciliary function, in the centrosomal region throughout all mitotic phases. Our work shows the dynamic role played by a TRIM family protein across various stages of mitosis for the first time, laying the foundation for exploring the therapeutic potential of TRIM8 in addressing cell cycle-related diseases, including cancer.

**HIGHLIGHTS:**

- TRIM8 is involved in transcriptional and post-translational regulation of “Cell Cycle Control of Chromosomal Replication” pathway and oversees the expression of *TOP2A*, essential for mitotic chromosome structure maintenance.
- The silencing of *TRIM8* induces changes in cellular translation activity and alters the expression pattern of key translational proteins. Additionally, *TRIM8*-silencing leads to an elevation of long non-coding RNA (lncRNA) *MALAT1* in the polysome-bound fraction.
- TRIM8 is identified as a novel ciliary protein.
- Silencing of *TRIM8* results in the upregulation of the centrosomal protein CEP170, and both proteins co-localize in the centrosomal region throughout all stages of mitosis.

**GRAPHICAL ABSTRACT:** 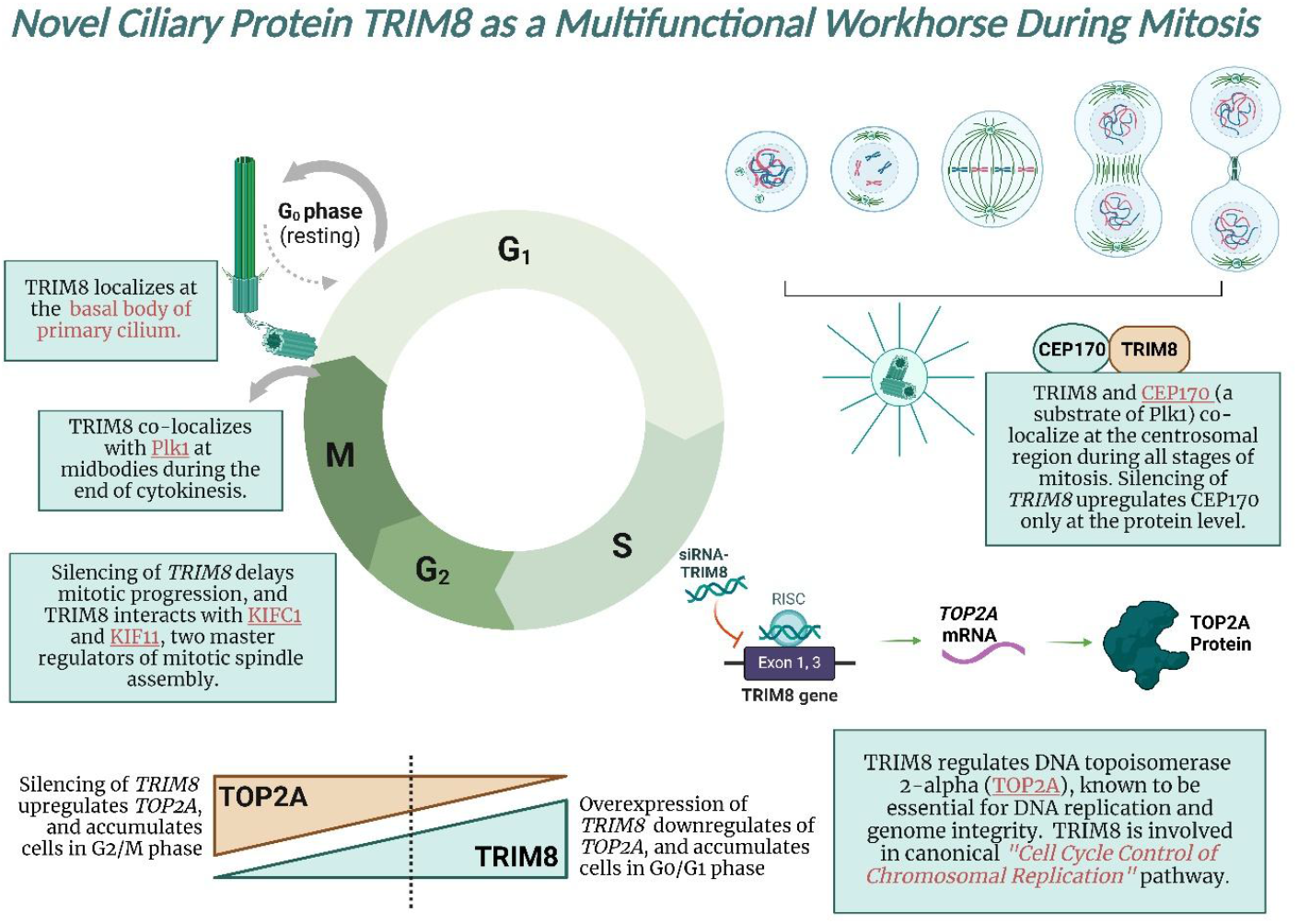

## 1. INTRODUCTION

Mitosis is one of the characteristic signatures of eukaryotic life and a fundamental process that plays a crucial role in growth & development, genetic stability, and tissue regeneration^1^. In eukaryotic organisms, the advancement of a cell throughout the mitotic cell cycle is highly regulated by different cyclin–CDK complexes^2–5^. Cyclin expression during the cell cycle is periodic due to a constant synthetic rate coupled with specific proteolysis mediated by the ubiquitin-proteasome system (UPS) or simply ubiquitination^6^. Indeed, ubiquitination plays a central role in regulating eukaryotic cell cycle transitions and checkpoints^6,7^, and it is a three-step process catalysed by E1 ubiquitin-activating enzymes, E2 ubiquitin-conjugating enzymes, and E3 ubiquitin–ligase enzymes^8^.

TRIM8 belongs to the **<u>TRI</u>**partite **<u>M</u>**otif (TRIM) family of E3 ubiquitin ligases that are known to share a structural similarity by having three common motifs or domains - one RING domain, one or two B-boxes, and a coiled-coil domain^9,10^. Previously, we termed TRIM8 as a “Molecule of Duality”, for its capacity to play a dual role in multiple contexts^11^. Notably, it can exert both anti-proliferative and pro-oncogenic activities, and operates in a manner that is not solely dependent on its E3 ligase function, showcasing both E3 ligase-dependent and -independent activities. Additionally, TRIM8 displays a broad subcellular influence, functioning in both the nucleus and cytoplasm to regulate NF-κB signaling. This multifaceted nature underscores TRIM8’s significance as a dynamic player in cellular processes with diverse and context-dependent impacts^11^. Although TRIMs are commonly known for their E3 ligase activity, many of these TRIMs can also function as transcriptional regulator, RNA binding protein (RBP) and can regulate autophagy^10,12^. Interestingly, quite a few numbers of TRIMs are shown to regulate TP53 expression, either by directly binding or indirectly by modulating upstream TP53 regulators^13^. And not surprisingly, the depletion of many TRIM proteins commonly leads to a significant cell cycle effect, marked by an elevated proportion of cells in the G0/G1 phase and a reduced fraction of cells in the S or G2-M phases. Specifically, the silencing of *TRIM14, TRIM27, TRIM28, TRIM29, TRIM52, TRIM59, TRIM66*, and *TRIM68* induces cell cycle arrest^14^. Earlier, we reported that TRIM8 physically interacts with KIF11 and KIFC1, two major regulators of mitotic spindle assembly formation and silencing of *TRIM8* results in an accumulation of cells in prometaphase that delays mitotic progression and in an increase of aneuploid cells^15^. Overall, there is a preliminary indication that TRIM8 may play a significant role in the mitotic cell cycle^15–17^; however, to date, no study has thoroughly investigated the involvement of TRIM8 during different stages of mitotic cell cycle progression and regulation. Indeed, there is no single descriptive study that elucidates the involvement of any TRIM family protein during different stages of mitosis and at various levels of mitotic cell cycle control in a systematic approach.

In this study, leveraging differential transcriptomic (scRNA-seq), translatomic (polysome RNA-seq), and proteomic (LC-MS/MS) approaches we aim to unravel the intricate role of TRIM8 across multiple levels—transcriptional, translational, and post-translational—and throughout distinct phases of the cell cycle (G0/G1, S, G2/M). We demonstrate that silencing *TRIM8* leads to the upregulation of both mRNA and protein expression of Topoisomerase IIα (TOP2A), crucial for chromosomal replication and compaction^18,19^, and upregulates the protein-level expression of five essential components of the minichromosome maintenance (MCM) protein complex, namely MCM3-7 helicases. An earlier study demonstrated that lncRNA *MALAT1* impedes *TOP2A* mRNA degradation, and the knockdown of *MALAT1* results in the downregulation of TOP2A^20^. Our polysome RNA-seq data shows an elevation of lncRNA *MALAT1* bound in polysome fractions upon silencing of *TRIM8* and unveils a new prospect for global translation control during the progression of mitotic cell cycle. Moreover, we identify TRIM8 as a novel ciliary protein. We show that *TRIM8* co-localises with the centrosomal protein CEP170, required to support ciliary function^21,22^, in the centrosomal region during all phases of mitosis. Our research uncovers how a TRIM family protein is involved in different stages of cell division, shedding light on its role from the beginning to the end of mitosis.

## 2. RESULTS

### 2.1. TRIM8 regulates the canonical “Cell Cycle Control of Chromosomal Replication” pathway, exerting negative control on TOP2A

Previously, we observed that *TRIM8*-silenced cells contribute significantly to an aneuploidy phenotype^15^, however, the underlying molecular mechanisms driving TRIM8’s involvement in chromosomal replication remain completely unknown. In this study, employing label-free differential proteomic analysis with liquid chromatography-tandem mass spectrometry (LC-MS/MS) upon synthetic siRNA-mediated silencing of *TRIM8* in hTERT-immortalized retinal pigment epithelial (hTERT RPE-1; henceforth referred to as RPE) cells, we identified 163 differentially expressed (DE) proteins (Supplementary Table 1). The activation *z-score* analysis with all DE proteins on QIAGEN Ingenuity Pathway Analysis (IPA) for canonical pathways revealed a positive *z-score* of 2.449 for the “Cell Cycle Control of Chromosomal Replication” pathway, and all DE proteins within this pathway were observed to be upregulated (Figure 1A). An interesting observation was made in the analysis conducted using enrichGO.R, focusing on cellular components, revealing upregulated proteins are primarily associated with cell cycle-related complexes, including chromosomal region, spindles, microtubules, DNA replication preinitiation complex, and MCM complex (Figure 1D). Conversely, downregulated proteins exhibit distinct associations, notably with components such as ribosomes, peptidase complexes, and proteasome complexes (Figure 1C). A parallel trend is also evident in the analysis of topmost significant QIAGEN IPA canonical pathways. The upregulated proteins predominantly align with cell cycle-associated pathways, while the downregulated proteins primarily impact immunity- and protein ubiquitination-related pathways (Figure 1E). The canonical “Cell Cycle Control of Chromosomal Replication” pathway also emerged as the top hit in the list of most enriched pathways when analysed with only upregulated proteins (Figure 1E) and exhibited a positive *z-score* with significant increase in the activation of biological function (Supplementary Figure 1). Notably, differential expression of TOP2A, vital for chromosomal replication, genomic integrity, and maintenance of mitotic genome structure^18,19^, and five key minichromosome maintenance (MCM) complex proteins (MCM3, MCM4, MCM5, MCM6, MCM7), point to TRIM8’s involvement in the canonical “*Cell Cycle Control of Chromosomal Replication*” pathway (Figure 2A and Supplementary Table 2). The elevated expression of TOP2A and five major MCM complex proteins in the “*Cell Cycle Control of Chromosomal Replication*” pathway (Figure 2B), underscores TRIM8’s crucial role in regulating components associated with chromosomal replication. This emphasizes TRIM8’s impact on the cell cycle, particularly in influencing DNA replication processes through the modulation of key regulatory proteins of chromosomal replication like TOP2A and members of the MCM complex.

**Figure 1:**
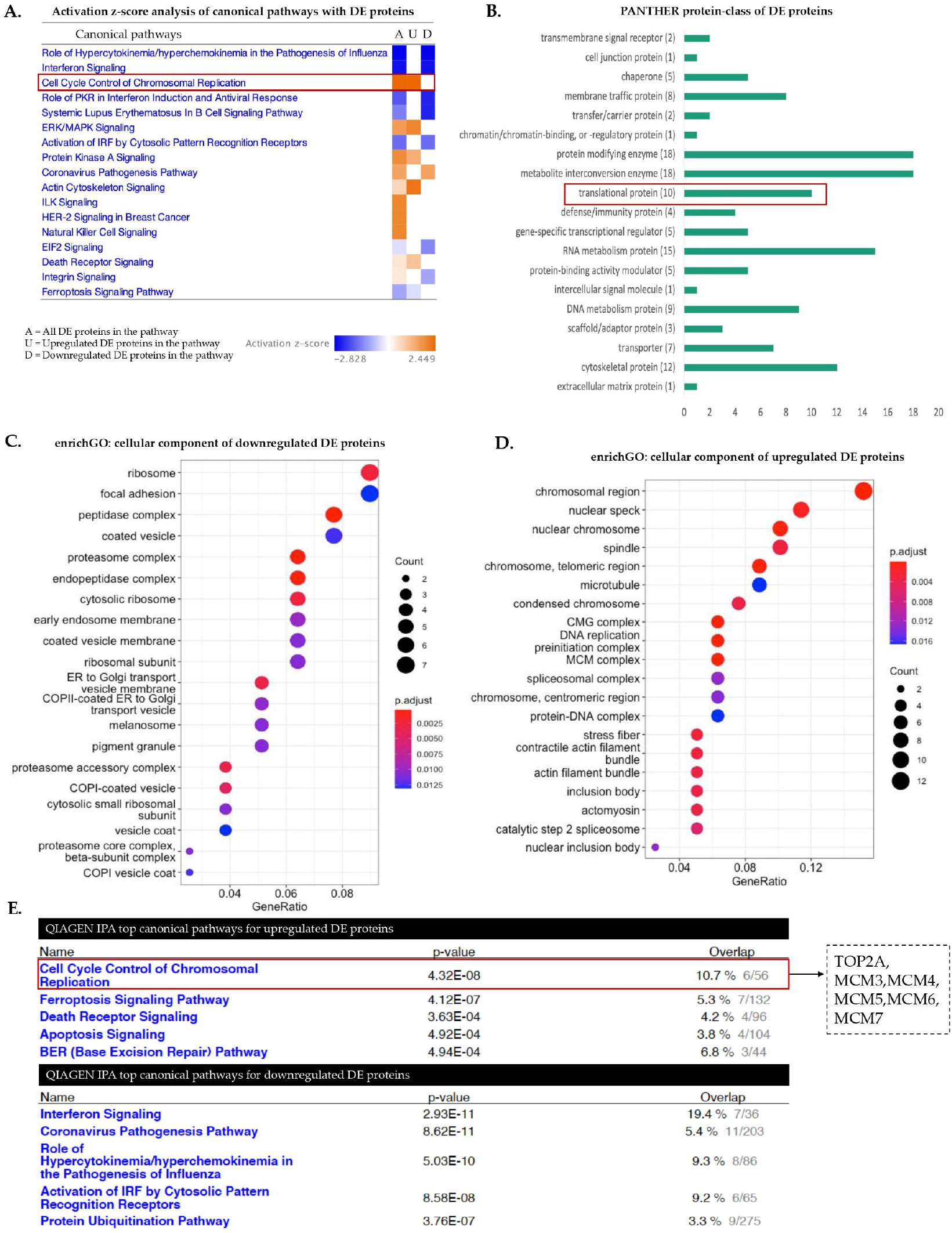
Involvement of TRIM8 in the “Cell Cycle Control of Chromosomal Replication” pathway. **A)** Activation z-score analysis conducted on QIAGEN IPA reveals that all differentially expressed (DE) proteins within the canonical “Cell Cycle Control of Chromosomal Replication” pathway from differential proteomic (LC-MS/MS) study in RPE cells exhibit upregulation upon *TRIM8* silencing. **B)** PANTHER analysis is utilized for the identification of protein classes among the DE proteins. **C) & D)** enrichGO analysis delineates the cellular components associated with both up- and down-regulated DE proteins. **E)** Top five pathways for up- and down-regulated DE proteins are separately identified using QIAGEN IPA.

**Figure 2:**
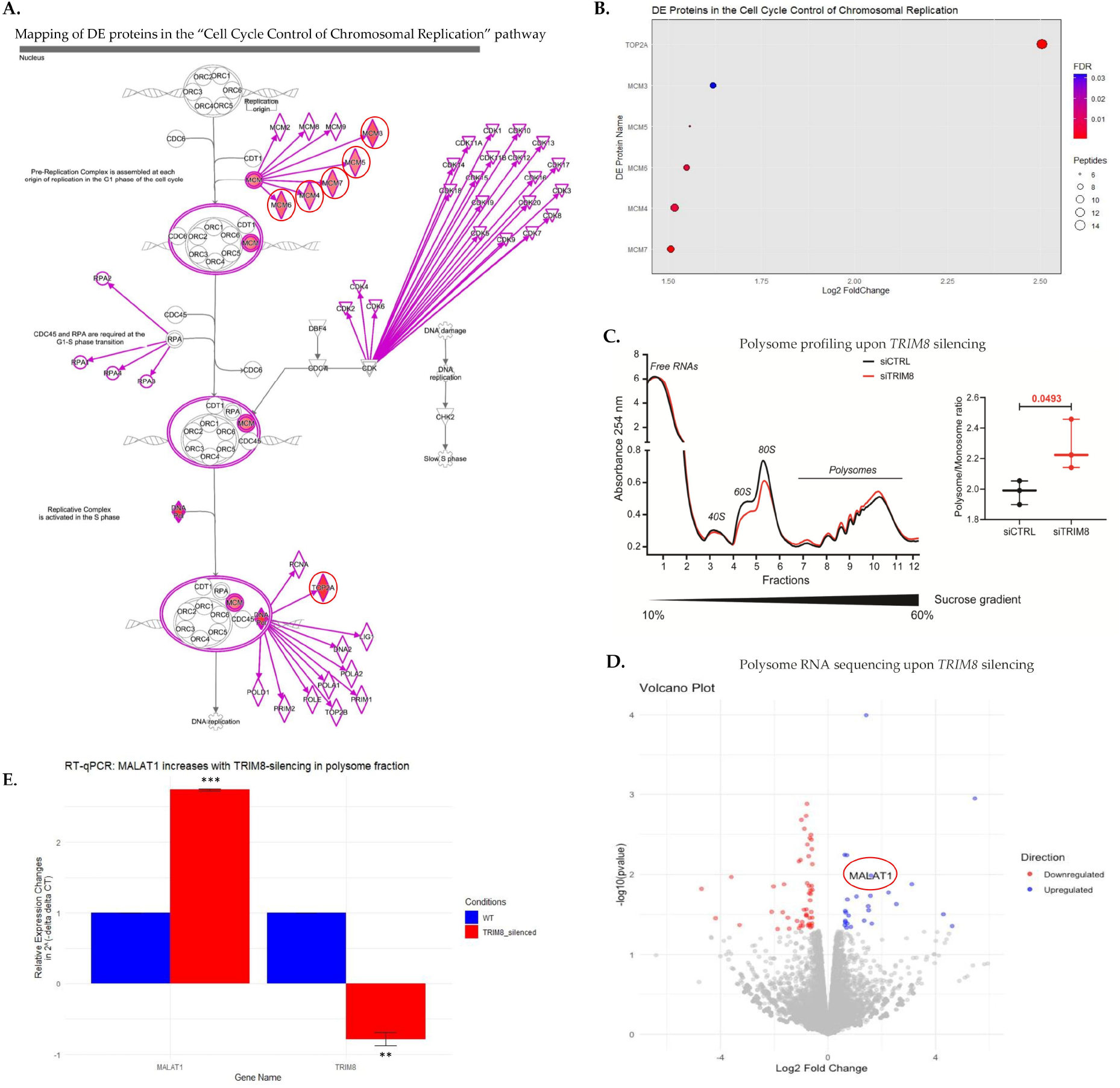
Differential proteomic (LC-MS/MS) and translatomic (polysome profiling with RNA-seq) study upon silencing of *TRIM8* in RPE cells. **A)** The positioning of differentially expressed (DE) proteins identified from the LC-MS/MS study within the canonical “Cell Cycle Control of Chromosomal Replication” pathway highlights TRIM8’s involvement in DNA replication, particularly through the regulation of TOP2A and MCM complex proteins (MCM3-7). DE proteins observed upon TRIM8 silencing are highlighted and encircled in red. **B)** Among the DE proteins, TOP2A emerges as the most significant and differentially expressed protein following TRIM8 silencing in the “Cell Cycle Control of Chromosomal Replication” pathway. **C)** Polysome profiling using sucrose density gradient centrifugation reveals alterations in translation activity in RPE cells upon *TRIM8* silencing, indicating its role in modulating translation. **D)** A volcano plot illustrates differentially translated genes, characterized by a |log2FC| ≥ 1.5 and a p-value < 0.05, identified from polysome RNA-seq upon *TRIM8* silencing in RPE cells. Notably, the long non-coding RNA *MALAT1* is upregulated in the polysome fraction sucrose density gradient centrifugation upon *TRIM8* silencing. **E)** RT-qPCR validation confirms the upregulation of *MALAT1* long non-coding RNA, validating its status as a differentially translated gene identified from polysome RNA-seq upon *TRIM8* silencing.

### 2.2. TRIM8 silencing modulates translational activity, eliciting an upregulation of polysome-bound MALAT1 lncRNA

A further protein-class analysis with PANTHER18.0 of the 163 differentially expressed proteins from LC-MS/MS study revealed that TRIM8 downregulation had a discernible impact on translational regulation, influencing ten important translational proteins (Figure 1B). Notably, seven translational proteins were upregulated, including RPS16, WARS, RPS9, CARS, RPL18A, ETF1, and RPL27A, while three were downregulated, including PABPN1, LRRC47, and RSL1D1 (Supplementary Table 1). Analysis of cellular components of all differentially expressed proteins with enrichGO.R showed “Ribosome” and “Cytosolic Ribosome” as one among the top and significant cellular components (Supplementary Figure 2). To further elucidate the effect of *TRIM8* downregulation on translational activity, polysome fractionations experiment on sucrose gradients was conducted, revealing a significant increase in the polysome to monosome ratio upon *TRIM8* silencing in RPE cells. This indicates an augmentation in translational activity when TRIM8 is perturbed (Figure 2C). In many previous studies, *MALAT1* has emerged among the long noncoding RNAs that play important role in cell cycle regulation^20,23^. Notably, *MALAT1* can act as a “sponge” for miR-561, impeding TOP2A mRNA degradation as a target of *miR-561* and knockdown of *MALAT1* has been associated with downregulation of TOP2A^20^. Interestingly, prior ribosome footprinting studies have also detected the presence of *MALAT1* in polysome fractions^24,25^. Here, our polysome profiling coupled with RNA-sequencing (RNA-seq) upon silencing of *TRIM8* unveiled an intriguing elevation of the long noncoding RNA *MALAT1* in the polysome-bound fraction (Figure 2D-E), suggesting a potential link between TRIM8, *MALAT1*, and translational modulation. This observation adds a layer of complexity to the potential connections between TRIM8, *MALAT1*, and translational modulation, shedding light on novel aspects of their interactions in cell cycle regulation. While detailed investigation was beyond the scope of this study, additional lncRNAs, such as *DOCK9-AS2* and *RNR2*, also exhibited perturbations in the polysome RNA-seq data following *TRIM8* silencing (Supplementary Table 3).

### 2.3. scRNA-seq unveils TRIM8’s dynamic role across mitotic phases (G0/G1, S, G2/M), and its regulation of TOP2A at the transcriptional level

To comprehend the impact of *TRIM8* downregulation at transcriptional level, BD Rhapsody™ single-cell RNA sequencing (scRNA-seq) and Whole Transcriptome Analysis (WTA) was conducted with an unsynchronized pool of RPE cells. Principal Component Analysis (PCA) conducted on the scRNA-seq data revealed two distinct components for the 2378 Wild Type and 2581 *TRIM8*-silenced cells, respectively, showing the clear impact of *TRIM8-*silencing on the data with highly dispersed genes (Figure 3A-B). As many of the cell cycle marker genes like *TOP2A* can vary in their expression in a population of cells that are at the different stages of cell cycle due to the heterogeneity of single-cell RNA-sequencing data^26,27^, so we tried to understand whether the transcriptional impacts of *TRIM8*-silencing on cell cycle marker genes like *TOP2A* are real or they are affected by the cellular heterogeneity. Recognizing the potential impact of cell cycle heterogeneity on such marker genes, we stratified cells into different cell cycle stages (G0/G1, S, G2/M) for a focused analysis to reveal the genuine impact of *TRIM8* downregulation on cell cycle-associated marker genes. PhenoGraph, a robust graph-based method for identifying subpopulations in high-dimensional single-cell data^28^, was applied to the total pool of cells using 95 cell cycle candidate marker genes (comprising 42 S-phase markers and 53 G2/M markers) from Tirosh et al. (2016)^27^. This analysis unveiled ten distinct clusters, as visualized on the two-dimensional tSNE space (Figure 3C). Subsequent gene expression pattern analysis of 95 cell cycle candidate marker genes among these ten clusters, conducted with the iCellR algorithm, further delineated three distinct clusters: cluster 1, cluster 2, and cluster 6, which exhibited specificity to G0/G1, S, and G2/M phases, respectively, based on the gene expression patterns of cell cycle marker genes (Figure 3D; Supplementary Figure 3). For instance, Cluster 6 (henceforth referred to as G2/M cluster) exhibited the highest number of differentially expressed G2/M marker genes, suggesting the predominance of cells in the G2/M phase within this cluster (Supplementary Figure 3). Each of the three cellular clusters was subsequently divided into wild type and *TRIM8*-silenced subgroups based on the Illumina sample tags 05 and 06, which were utilized during library preparation for siRNA negative controls and TRIM8-siRNA, respectively. Within each cluster, differentially expressed (DE) genes were subsequently identified between the *TRIM8*-silenced and control RPE populations of cells using the SeqGeq™ 1.7.0 bioinformatics analysis platform, tailored for BD Rhapsody scRNA-seq data analysis. Identification of DE genes at G2/M cluster (Figure E-F) showed that *TOP2A*, that itself is one of the G2/M marker genes, was indeed differentially expressed upon silencing of *TRIM8* in G2/M cluster (Figure 3G-H). This data showed the real transcriptional elevation of *TOP2A* upon silencing of *TRIM8* when compared between *TRIM8*-silenced cells over control RPE cells at the G2/M phase (Figure 3G-H). Additionally, stage-specific analysis indicated that *TOP2A* is affected across all three clusters i.e., G0/G1, S, G2/M (Figure H-J). Conducting QIAGEN IPA analysis with DE genes from each cluster unveiled enriched pathways specific to different phases of mitosis following *TRIM8* silencing (Figure 4A). Notably, in S-phase cells, all upregulated DE genes from *TRIM8*-downregulation enriched the “Cell Cycle: G1/S Checkpoint Regulation” pathway, while upregulated DE genes from G2/M phase cells showed enrichment in the “Kinetochore Metaphase Signaling” alongside “Cell Cycle Control of Chromosomal Replication” pathway (Figure 4A). These findings align with our previous studies demonstrating TRIM8’s interaction with kinetochore family proteins and its role in mitotic spindle assembly^15^. The data collectively emphasizes TRIM8’s regulatory impact on cell cycle processes during the progression of mitosis, particularly the transcriptional upregulation of *TOP2A*, a key member of the “Cell Cycle Control of Chromosomal Replication” pathway, that was differentially expressed in all three mitotic phases upon downregulation of *TRIM8*. Furthermore, SMC2 (Structural Maintenance Of Chromosomes 2), which exhibited upregulation in the differential proteomic analysis subsequent to *TRIM8*-silencing (see Supplementary Table 1), was observed to display a positive correlation with *TOP2A* mRNA expression across various cancers in The Cancer Genome Atlas (TCGA) dataset^29^, comprising 967 cancer samples sourced from the Cancer Cell Line Encyclopedia (Figure 3K). These findings also suggest a potential link between the upregulation of SMC2 and the transcriptional elevation of TOP2A upon *TRIM8*-silencing. Given that both SMC2 and TOP2A are integral components involved in the regulation of chromosomal structure maintenance and dynamics during various stages of the cell cycle^18,19,30–32^, their coordinated expression patterns underscore a putative role for TRIM8 in orchestrating processes crucial for chromosomal integrity and stability **(see Supplementary Methods for additional details)**.

**Figure 3:**
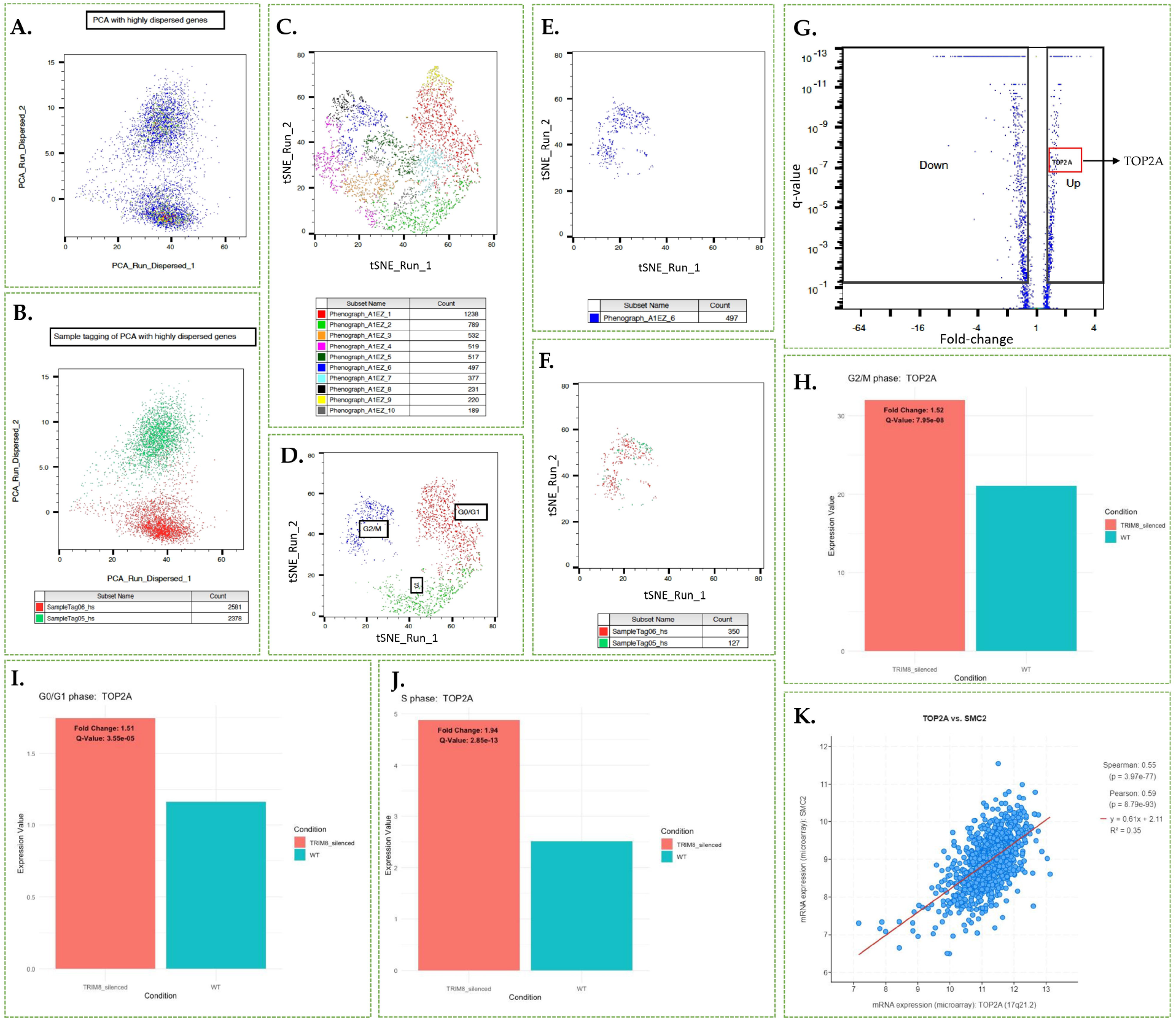
Impact of *TRIM8* silencing on single-Cell RNA-seq (scRNA-seq) analysis in unsynchronized RPE Cells. **A)** Dimensionality reduction was conducted using SeqGeq™ with Principal Component Analysis (PCA) machine learning algorithm on quality cells utilizing highly dispersed genes. **B)** PCA analysis clearly delineates two distinct components for wild-type (green) and TRIM8-silenced (red) RPE cells. Illumina SampleTag05 represents wild-type cells, while SampleTag06 represents *TRIM8*-silenced RPE cells. A total of 2378 quality cells were identified for the wild-type, and 2581 quality cells were identified for *TRIM8*-silenced RPE cells. **C)** Principal component parameters from scRNA-seq data, derived from highly dispersed genes, were mapped into two-dimensional tSNE space. Subsequently, PhenoGraph, a robust graph-based method for identifying subpopulations in high-dimensional single-cell data, was applied to the entire cell pool using 95 known cell cycle candidate marker genes. This process generated 10 PhenoGraph clusters of cells, effectively partitioning the high-parameter scRNA-seq data into phenotypically distinct subpopulations based on the expression pattern of 95 cell cycle marker genes. **D)** iCellR analysis on tSNE space was employed to discern the differential expression pattern of cell cycle marker genes among the clusters, identifying three major clusters: G0/G1, S, and G2/M. **E)** The G2/M cluster isolated on the tSNE space. **F)** SampleTag05 and SampleTag06 are utilized to distinguish wild-type and *TRIM8-*silenced cells within the G2/M cluster. **G)** Volcano plot depicts differential gene expression within the G2/M cluster, highlighting *TOP2A* as the upregulated gene during G2/M phase upon *TRIM8* silencing. **H), I), and J)** Bar plots demonstrate the upregulation of *TOP2A* in all three clusters or mitotic stages upon *TRIM8* silencing in the scRNA-seq data. **K)** The Cancer Genome Atlas (TCGA) dataset analysis, comprising 967 cancer samples from the Cancer Cell Line Encyclopedia, illustrates a moderate positive correlation in mRNA expression patterns between *TOP2A* and *SMC2*.

### 2.4. TRIM8 overexpression leads to cell cycle arrest at G0/G1 and downregulation of TOP2A

In our earlier study, *TRIM8* silencing led to an accumulation of cells at prometaphase^15^. In the current stage-specific scRNA-seq analysis, all upregulated DE genes from *TRIM8* silencing enriched the “Cell Cycle: G1/S Checkpoint Regulation” pathway in the S-phase cluster (Figure 4A). To investigate if TRIM8 overexpression could induce checkpoint regulation arrest, we conducted a flow cytometry study coupled with BrdU/7-AAD treatment assay in synchronized wild-type and *TRIM8-EGFP* transfected RPE cells (Figure 4B). Results showed that at the specified time-point, consistent with RPE cell standards (see materials and methods), while 89% of Wild-type RPE cells were in S-phase, a substantial proportion of *TRIM8-EGFP* transfected cells remained in the G1 window, unable to progress to S-phase (Figure 4C). Earlier report showed that *TOP2A* downregulation in JEG-3 cells resulted in G0/G1 phase arrest^33^. Hajibabaei, Sara et al. (2023) showed that knockdown of *MALAT1* also arrested mitotic progression at G0/G1 and downregulated *TOP2A* in breast cancer cells^20^. As our scRNA-seq study indicates that *TRIM8* silencing upregulates *TOP2A* and flow cytometry results show that *TRIM8* overexpression leads to G0/G1 phase arrest, we hypothesized whether *TRIM8* overexpression could result in *TOP2A* downregulation. RT-qPCR data upon *TRIM8-EGFP* overexpression in an unsynchronized RPE cell population revealed a significant downregulation of *TOP2A* mRNA (Figure 4D). This data establishes that TRIM8 negatively regulates *TOP2A*, influencing the “Cell Cycle Control of Chromosomal Replication” pathway, and concurrently participates in G1/S checkpoint regulation pathway possibly through transcriptional level control of *TOP2A*.

**Figure 4.**
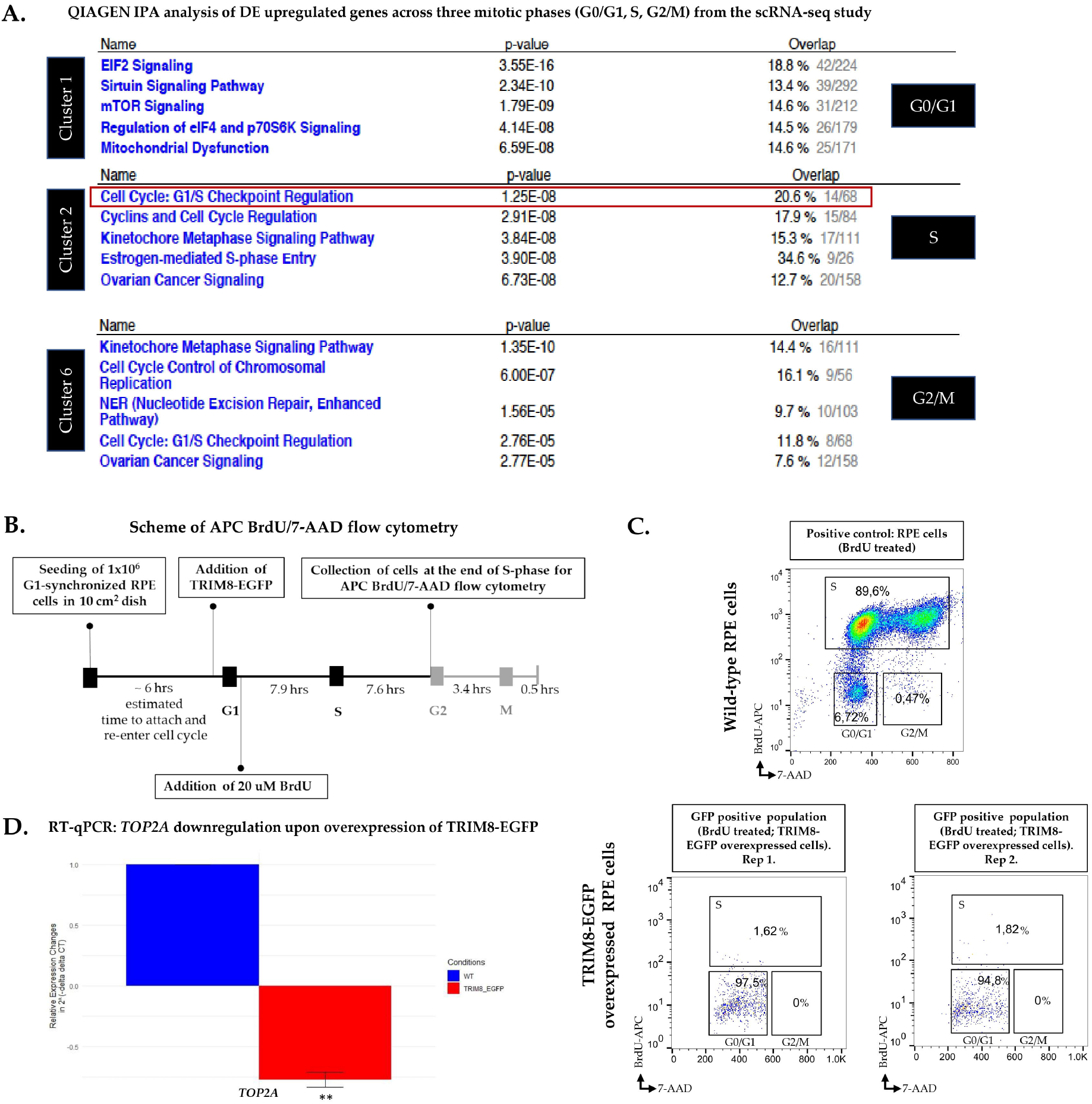
*TRIM8* overexpression leads to cell cycle arrest at G0/G1 and downregulation of *TOP2A* in RPE cells. **A)** QIAGEN Ingenuity Pathway Analysis (IPA) analysis of DE upregulated genes of three mitotic phases (G0/G1, S, G2/M) from the scRNA-seq data. Silencing of *TRIM8* perturbs “Cell Cycle: G1/S Checkpoint Regulation” pathway significantly at the Cluster 2 or S-phase. **B)** Schematic design of the APC BrdU/7-AAD assay for flow cytometry analysis, illustrating the time points of TRIM8-EGFP and BrdU treatments. **C)** RPE cells were first synchronized and collected at G1. G1 RPE cells were seeded again to restart cell cycle uniformly and treated with BrdU. At the end of the S-phase when only BrdU-labelled WT RPE cells were in S-phase, the BrdU-labelled TRIM8-EGFP overexpressed cells were found arrested at G0/G1 phase in the flow cytometry analysis. This shows overexpression of TRIM8-EGFP leads to G0/G1 arrest in synchronized RPE cells. **D)** RT-qPCR validation shows that overexpression of TRIM8-EGFP leads to downregulation of *TOP2A*.

### 2.5. TRIM8: a ciliary protein that posttranslationally regulates CEP170 and co-localizes with it across mitotic phases

The classical model of centrosome biogenesis indicates that perturbations in centrosomes of certain vertebrate cells result in G1/S arrest, attributed to the loss of centrosome integrity^34^. The centrosome consistently functions as a major activity centre during mitosis^35^ and in many cell types at the onset of G1/S transition, the centriole pair divides, initiating the growth of new centrioles from the periphery of the original two centrioles^36^. In our previous mass-spectrometry-based interactomic study, CEP170 was screened among fifty putative TRIM8-interacting proteins, including essential mitotic assembly proteins such as KIF11 and KIFC1^15^. CEP170 maintains constant expression throughout the cell cycle, and it serves as a discriminating marker between bona fide centriole overduplication and centriole amplification resulting from aborted cell division^37^. Additionally, CEP170 is a physiological substrate of Plk1 kinase and undergoes phosphorylation by Plk1^37^. Interestingly, our earlier study showed that TRIM8 also colocalizes with Plk1 at the midbodies during the end of cytokinesis^15^. Here, we overlapped 163 differentially expressed proteins identified from our differential proteomic (LC-MS/MS) study performed upon silencing of *TRIM8* (Supplementary Table 1) with the 582 Human Centrosomal Proteins retrieved from Human Protein Atlas (HPA). This reveals five common centrosomal proteins, CEP170, CKAP5, PXN, RBM39, and TAP1 as differentially expressed in our proteomic study that are perturbed upon silencing of *TRIM8* (Figure 5A). Although CEP170 was screened as an upregulated protein in our differential proteomic study (Supplementary Table 1), further examination of our scRNA-seq and polysome-RNA-seq data revealed no discernible perturbation of CEP170 at both the transcriptional and translational levels (data not shown). This substantiates the notion that TRIM8 possibly regulates CEP170 through post-translational mechanisms. In line with the findings from our differential proteomics study, an immunofluorescence study was conducted to explore the co-localization of TRIM8 with CEP170. The observations reveal that TRIM8 co-localizes with CEP170 at centrosomes throughout all phases of mitotic cell division in HeLa cells, spanning from the initial interphase to the conclusion of telophase. During interphase, TRIM8 has a tendency to exhibit a scattered nuclear dots distribution, yet maintains consistent colocalization with CEP170, particularly at the centrosomal region during all phases of mitosis (Figure 6A). Along with TRIM8 and CEP170, γ-tubulin, a marker of the centrosomal region, was also observed (Supplementary Figure 4). Guarguaglini et al. (2005) showed that during early interphase CEP170 localises at the mother centriole^37^, the site of primary cilium formation^38^. Zhang et al. (2019) showed that mechanistically, WDR62 promotes CEP170’s localization to the basal body of primary cilium^22^. A more recent work showed that CEP170 is required for supporting ciliary function^21^. Here, we performed an overlap study between 302 SYSCILIA Gold Standard Ciliary Proteins^39,40^ and our 163 differentially expressed proteins from LC-MS/MS analysis. We identified NUP93 and RANBP1 as two common differentially expressed gold-standard ciliary proteins upon silencing of *TRIM8* (Figure 5B-C). MCL (Markov Cluster Algorithm) clustering of the 163 differentially expressed proteins exposed a cluster that showed an enrichment in “Ciliary Landscape” (WikiPathways: WP4352) with FDR: 2.30e-17 and that also includes CEP170 along with MCM complex proteins (Figure 5D). As primary cilium formation occurs during G0/G1 phase and disassembles upon cell cycle re-entry^38,41,42^, we investigated our G0/G1 differentially expressed genes from the scRNA-seq analysis for ciliary gene enrichment. Analysis revealed an enrichment of SYSCILIA Gold Standard ciliary genes within this cluster, indicating their involvement in the biological process “Cilium Assembly” (Figure 5E). However, in publicly accessible databases such as CiliaCarta, only two TRIM E3 ubiquitin ligases, TRIM32 and TRIM59 are annotated as ciliary genes, whereas TRIM8 is classified as completely “unknown” with regards to its subcellular localization at the primary cilium^43,44^. Therefore, following our own computational analysis and prediction, as mentioned above, an immunofluorescence study was carried out to validate whether TRIM8 can localise at the primary cilium upon serum starvation that induces cell cycle arrest in the G0/G1 phase and ciliary assembly in RPE cells^45^. The results unequivocally demonstrate the localization of TRIM8 at the site of primary cilium formation (Figure 6B). Additionally, we showed that TRIM8 colocalizes with CEP170 during both starvation and non-starvation conditions (Supplementary Figure 5). Previous studies have identified CEP170 as a basal foot cap protein^46^. Given TRIM8’s localisation at the site of primary cilium formation (Figure 6B) and its co-localisation with CEP170 during both starvation and non-starvation conditions, we argue that TRIM8 is also a ciliary protein localising to the basal body region. Although TRIM8 exhibits a dispersed nuclear-dot localisation during interphase, its co-localisation with CEP170 remains consistent throughout all phases of mitosis in the centrosomal region (Figure 6A). For both the TRIM8-CEP170 co-localization study and the investigation of TRIM8 localization at the primary cilium, the findings were further validated in two additional cell lines, HEK293 and SH-SY5Y (Supplementary Figures 6 and 7).

**Figure 5.**
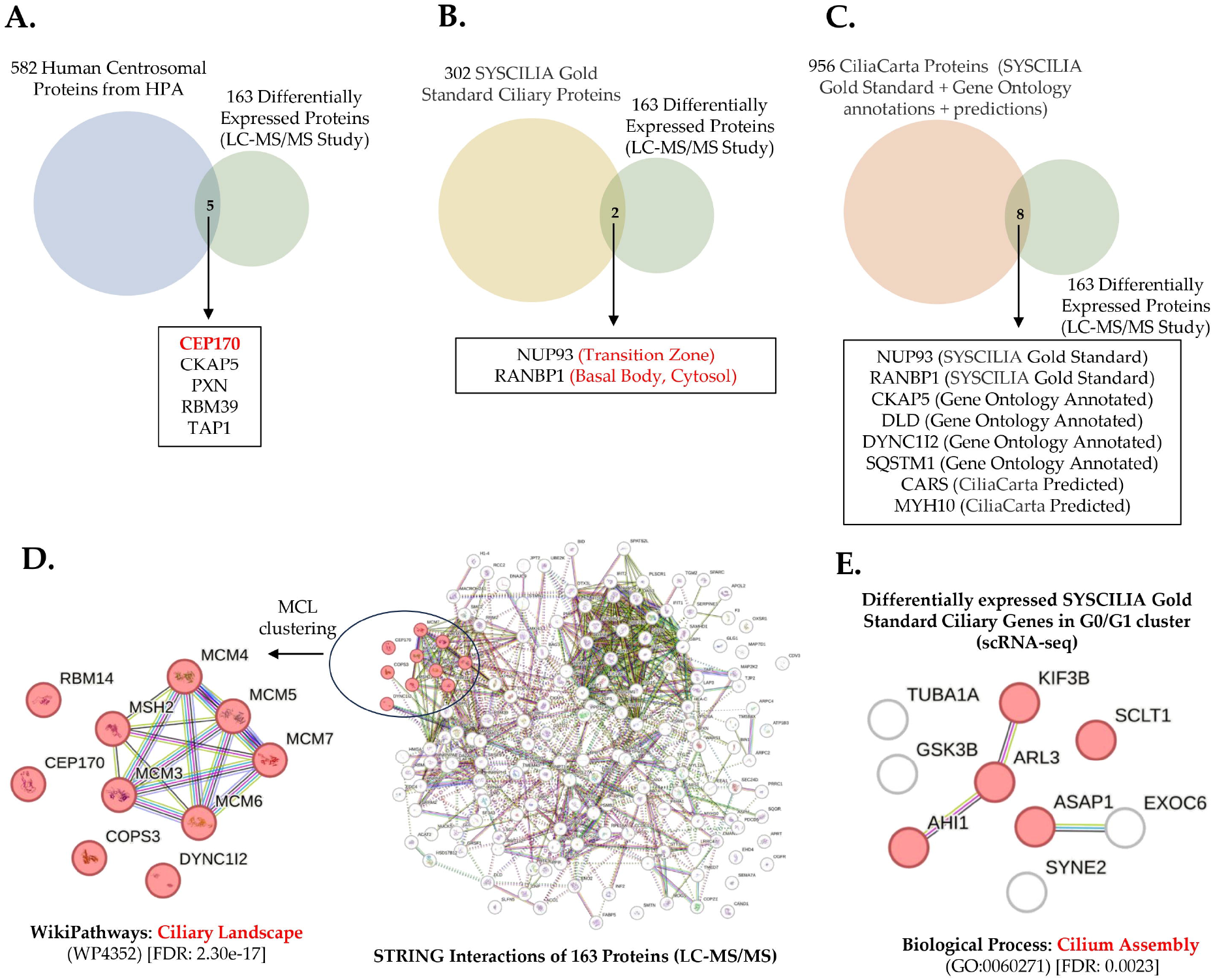
Network analysis to predict TRIM8’s involvement in primary cilium. **A)** Overlap between 582 human centrosomal proteins from Human Protein Atlas (HPA) and 163 DE proteins upon *TRIM8* silencing from the differential proteomic (LC-MS/MS) study. **B)** Overlap between 302 SYSCILIA Gold Standard Ciliary proteins and 163 DE proteins. **C)** Overlap between 956 CiliaCarta Proteins (combined with 302 SYSCILIA Gold Standard proteins and proteins from Gene Ontology annotations & predictions) and 163 DE proteins. **D)** STRING analysis of nine differentially expressed SYSCILIA Gold Standard ciliary genes in G0/G1 cluster identified from the scRNA-seq study. Five DE genes (KIF3B, ARL3, AHI1, SCLT1, and ASAP1) show to be involved in the biological process Cilium Assembly (GO:0060271). **E)** MCL graph clustering performed upon the STRING protein-protein interactions of 163 DE proteins from LC-MS/MS study identifies a unique cluster of ten DE proteins, including CEP170 and MCM complex proteins, that is enriched in ciliary landscape (WP4352) with FDR: 2.30e-17.

**Figure 6.**
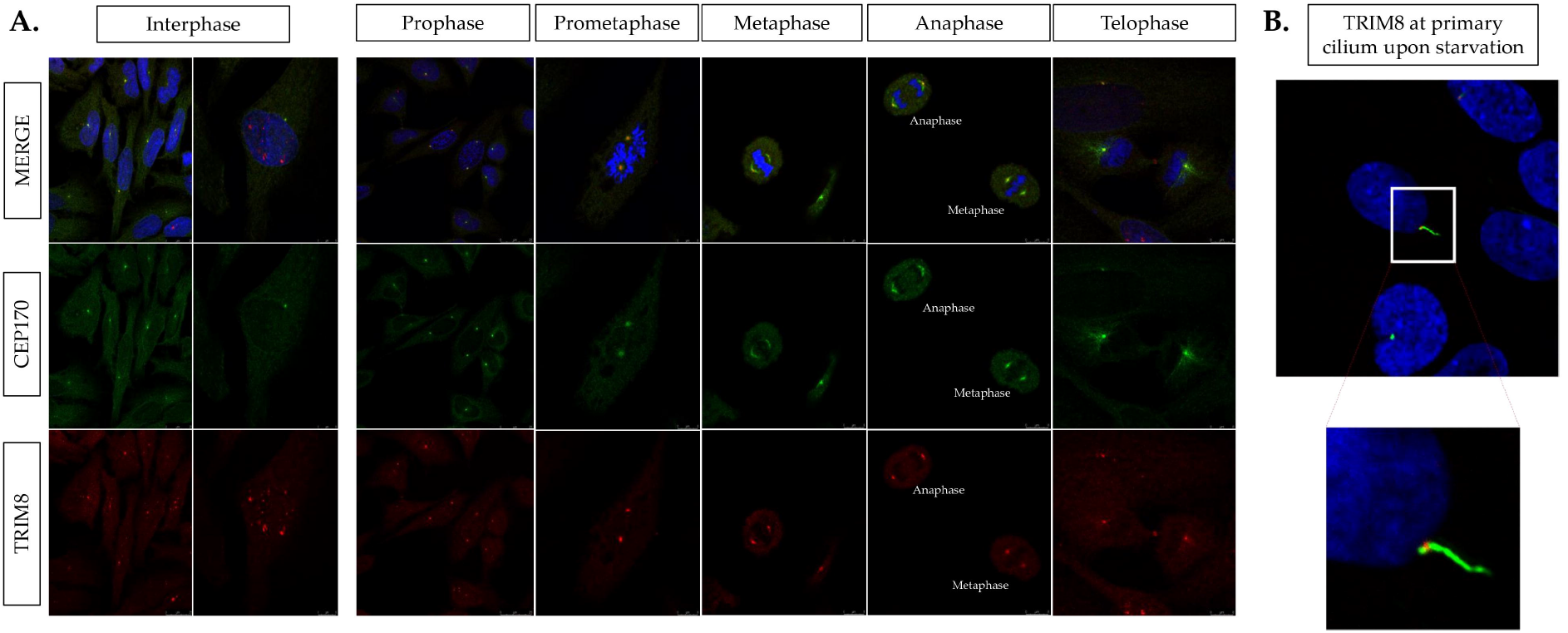
TRIM8 is a novel ciliary protein and co-localises with CEP170 in the centrosomal region during all mitotic phases. **A)** TRIM8 (red) co-localizes with CEP170 (green) in HeLa cells throughout all mitotic stages. Cell nuclei were stained with DAPI (blue). **B)** TRIM8 (red) localizes at the site of formation of the primary cilium, induced to form upon serum starvation in RPE cells. The primary cilium (green) was identified using anti-acetylated alpha tubulin. **…**

## 3. DISCUSSION

In our study, leveraging existing algorithms, we optimised an approach to analyse single-cell transcriptomic data from a “single cell line”, without mitigating the effect of cell-cycle heterogeneity. This enabled us to discern the genuine impact of *TRIM8* downregulation on mitotic cell cycle candidate genes without the confounding effects of cell-cycle variability of scRNA-seq data generated from unsynchronized population of cells from a single cell line. Our comprehensive investigation with a multiomics strategy elucidated the involvement of *TRIM8* in different levels (transcription, translation, post-translation) and stages (G0/G1, S, G2/M) of cell cycle regulation & progression in a unique way and highlighted the intricate connection between TRIM8, TOP2A, and MCM complex proteins during chromosomal replication. The observed upregulation of TOP2A and MCM proteins upon *TRIM8* silencing provides novel insights into TRIM8’s role in chromosomal replication and stability. Although in our earlier work we reported that silencing of *TRIM8* results in aneuploidy phenotype^15^, this study provides the first to offer foundational and conceptual molecular insights into the involvement of TRIM8 in chromosomal replication pathway through regulation of TOP2A, known to be essential for chromosomal replication and maintenance of mitotic chromosome structure^18,19,32,33^. Another insightful observation made in this is a notable upregulation of polysome-bound *MALAT1* lncRNA upon *TRIM8* downregulation, as observed in polysome RNA-seq analysis. Previous findings indicate that *MALAT1* depletion induces G0/G1 phase cell cycle arrest^20^. In this study, we showed *TRIM8* overexpression also led to cell accumulation at the G0/G1 phase. Notably, *MALAT1* has been linked to miR-561 “sponging,” inhibiting *TOP2A* mRNA degradation, where *MALAT1* knockdown resulted in *TOP2A* downregulation^20^. Earlier studies reported *MALAT1*’s presence in polysome fractions^24,25^, and influence on RNA translation^47,48^. Intriguingly, our polysome RNA-seq data shows significantly higher *MALAT1* lncRNA bound in polysome fractions, while RT-qPCR, scRNA-seq, and LC-MS/MS data demonstrate *TOP2A* upregulation upon *TRIM8* silencing. This introduces the possibility of global translation control during mitotic cell cycle, warranting in-depth investigation, particularly focusing on the TRIM8-*MALAT1*-TOP2A axis. The recent surge in interest surrounding Liquid-Liquid Phase Separation (LLPS) has provided insights into precise spatial and temporal regulation within living cells^49,50^. LLPS, forming biomolecular condensates similar to earlier-known membrane-less organelles (MLO), segregates nucleic acids and proteins into micron-scale liquid-like bodies^51^. TOP2A, identified as an LLPS “client,” associates with specific biomolecular condensates, including the nucleolus, centrosome, spindle pole body, and P-body^52,53^. Additionally, MCM complex proteins, particularly MCM4-7, function as “regulators” in the same biomolecular condensate categories, such as “centrosome/spindle pole body”^52,53^. Although we don’t have any empirical evidence to prove the hypothesis, the localization of TRIM8 at the centrosomal region throughout all mitotic phases and the impact of *TRIM8* silencing on TOP2A and MCM complex proteins necessitate future mechanistic studies to establish the functional relationship of the TRIM8-TOP2A-MCM axis. There is a significant possibility that the “once per cell cycle” mechanism, associated with both chromosomal replication and centrosomal duplication, has precursors at the centrosome/spindle pole body condensate—effectively remaining one of the most active centers of mitosis throughout the cell cycle.

Interestingly, potential functional studies and future work on TRIM8 at the centrosomal region could draw inspiration from a recent report establishing the role of another TRIM E3 ubiquitin ligase family protein, TRIM37, with polo-like kinase 4 (Plk4), and a centrosomal protein, CEP192^54^. TRIM37 can cause a significant reduction in the centrosomal protein CEP192 when overexpressed in RPE-1 cells. This overexpression also impedes the assembly of acentrosomal spindles, resulting in mitotic failure. Conversely, the absence of TRIM37 improves cell proliferation and facilitates acentrosomal spindle assembly^54^. The essential kinase Plk4, crucial for centriole duplication, relies on recruitment to the centrosome, facilitated by centrosomal proteins CEP192 and CEP152^54–56^. Recent findings propose that TRIM37 plays a role in regulating CEP192 stability, and in preventing self-assembly of Plk4 to form condensates that nucleate microtubules rather than controlling the expression of Plk4^54^. In summary, TRIM37 operates bidirectionally: low TRIM37 levels support the formation of robust acentrosomal spindle assembly by allowing Plk4 to form condensates that nucleate microtubules, while high TRIM37 levels lead to CEP192 degradation and no Plk4 condensates, causing acentrosomal spindle assembly failure^54^. Translating these observations to our study on TRIM8, we identify three crucial points within the axis of TRIM8, polo-like kinase 1 (Plk1), and CEP170. Firstly, *TRIM8* silencing upregulates CEP170, akin to TRIM37’s impact on CEP192 at the protein level. Secondly, although we reported earlier that TRIM8 and Plk1 colocalize at midbodies during the end of cytokinesis^15^, in our current differential transcriptomic (scRNA-seq) and proteomic (LC-MS/MS) study we could not observe any effect on Plk1 expression upon *TRIM8* silencing (data not shown), mirroring TRIM37’s no role in controlling Plk4 expression. Thirdly, TRIM8 and CEP170 exhibit centrosomal co-localization throughout mitosis, emphasizing their potential collaboration. In a previous study, CEP170 was found to localize at the mother centriole during early interphase^37^, now recognized as the site of primary cilium formation^38^. Remarkably, our immunofluorescence study showed that TRIM8 localises at the site of primary cilium formation and silencing of *TRIM8* perturbs NUP93, RANBP1, two gold-standard ciliary proteins. This study represents the first evidence identifying TRIM8 as a novel ciliary protein. Additionally, our mass-spectrometry (LC-MS/MS) analysis identified a distinct cluster of differentially expressed proteins using the MCL algorithm. Notably, this specific cluster demonstrates enrichment in ciliary landscape, featuring key proteins such as CEP170 and components of the MCM complex. This is consistent with the recent theory proposing that MCM-complex components not only localize to centrosomes but also play an independent role in ciliogenesis^57,58^. Altogether, our study propounds that TRIM8 is a multifunctional workhorse during mitosis and probably participate in a conserved “once per cell cycle” mechanism that is maintained from centrosomal duplication to chromosomal replication to the end of cytokinesis^59,60^. Recent studies, including those on glioblastoma, have highlighted the dual role of TRIM8 as both a tumor suppressor and oncogene. This complexity suggests that TRIM8-based therapies must carefully balance its functions depending on the specific cancer context^11,61–64^. For instance, targeted modulation of TRIM8’s interactions with mitotic regulators such as TOP2A and MCM complexes could provide a strategy for precise cancer treatment, where its tumor-suppressing effects are harnessed without exacerbating its oncogenic potential. Additionally, while our findings provide insights into TRIM8’s role in mitosis, further research using diverse cell models and advanced imaging techniques is needed to fully elucidate its mechanistic involvement. One limitation of this study is its reliance on a single-cell line model. Although the RPE cell line is a well-established model system for mitotic cell studies, it may not capture the full complexity of TRIM8 function across different cellular contexts in human tissues or primary patient-derived models. Addressing these gaps could deepen our understanding of TRIM8’s role in mitotic regulation and its therapeutic potential.

## 4. MATERIALS AND METHODS

### 4.1. Cell culture

hTERT-immortalized retinal pigment epithelial cells (hTERT RPE-1; hereafter RPE) and HeLa cells were cultured in Dulbecco’s Modified Eagle Medium (DMEM) supplemented with 4.5 g/L glucose, UltraGlutamine™ with sodium pyruvate (BE12-604F/U1), 10% filtered fetal bovine serum (FBS), and 1% penicillin-streptomycin mixture (DE17 602F).

### 4.2. siRNA-mediated silencing of TRIM8

A mix of Stealth siRNAs (Thermo Fisher Scientific; HSS129955, HSS129956, HSS188606) targeting exons 1 and 3 of *TRIM8* gene and a Stealth RNAi™ siRNA Negative Control for background comparison were transfected following the Lipofectamine® RNAiMAX Transfection protocol (Invitrogen™). 6×10^5^ RPE cells were seeded onto a 10 cm dish. Lipofectamine® RNAiMAX Transfection Reagent was diluted in Opti-MEM™ Reduced-Serum Medium according to the manufacturer’s instructions. The transfection reagent was incubated with the cells for 24 hours, and the final concentration of *TRIM8*-siRNA mix used was 80 pmols for each 10 cm dish. After 24 hours of transfection, RPE cells were harvested for subsequent experimental analyses.

### 4.3. RNA isolation, cDNA preparation, and quantitative reverse transcription PCR (RT qPCR)

Total RNA from RPE cells transfected with a *TRIM8*-siRNA or with the siRNA negative control was isolated as per the instruction of QIAGEN RNeasy Mini Kit for purification of total RNA. Upon isolation, reverse transcription was done using the Quantitect Transcription kit (Qiagen), according to the manufacturer’s instructions. DNA oligonucleotide primers for RT-qPCR were designed using the Applied Biosystems™ Primer Express™ software with default parameters. GAPDH and EEF1A1 were used as reference genes in the experiment. The reactions were run in triplicate in 10 ul of final volume with 10 ng of sample cDNA, 0.3mM of each primer, and 1XPower SYBR Green PCR Master Mix (Thermo Fisher Scientific-Applied Biosystems). Reactions were set up in a MicroAmp™ Optical 384-Well Reaction Plate with Barcode and run in an ABI Prism7900HT (Thermo Fisher Scientific-Applied Biosystems) with default amplification conditions. Raw Ct values were taken from SDS 2.4 (Applied Biosystems). Normalisation and relative changes in gene expression were quantified using 2^−ΔΔCt^ method.

### 4.4. Differential proteomic study (LC-MS/MS)

A label-free differential proteomic investigation employing liquid chromatography-tandem mass spectrometry (LC-MS/MS) was undertaken to elucidate the impact of TRIM8 gene silencing on retinal pigment epithelial (RPE) cells. Three replicates of RPE cells were transfected either with TRIM8-siRNA or a negative control siRNA. Transfected cells were harvested and subjected to lysis with 5% SDS and 50 mM ammonium bicarbonate. The lysed cells underwent three rounds of sonication on ice for 10 seconds each to ensure efficient membrane disruption. Following centrifugation at 13,000 rpm at 4°C, protein extracts were meticulously collected. A 50 μg portion of protein extract from each sample was enzymatically digested with trypsin using S-trap cartridges (ProtiFi), following the manufacturer’s protocol. The resulting peptide mixtures were then analyzed via nano LC–MS/MS by using an Easy-nLC II chromatographic system coupled with a linear trap quadrupole (LTQ) Orbitrap XL mass spectrometer (ThermoFisher Scientific, Waltham, MA). Chromatographic separation was achieved by fractionating samples onto a C18 capillary reverse-phase column (200 mm length, 75 μm inner diameter, 5 μm particle size) at a flow rate of 250 nL/min, employing a linear gradient of eluent B (0.2% formic acid in 95% acetonitrile) and eluent A (0.2% formic acid in 2% acetonitrile) from 5% B to 50% B over 260 minutes. Tandem mass spectrometric (MS/MS) analyses were performed in data-dependent acquisition mode, scanning the mass range from 400 to 1800 m/z. The top 10 most abundant ions in each MS scan were selected for MS/MS scans, with a dynamic exclusion window of 40 seconds to prevent repeated ion selection. Output files from the mass spectrometer were analyzed using MaxQuant software for peptide identification and quantification. Proteins were identified based on a minimum of 4 peptides, with at least 2 peptides being unique. False discovery rates (FDR) for reliable peptide spectrum match (PSM) and protein identification cutoffs were set to 0.01 to ensure high-confidence protein identification. Statistically significant proteins were evaluated using Perseus software, setting 5% FDR as the cutoff for the Student’s t-test. Fold Changes (FCs) were calculated by dividing the mean of significant proteins LFQ intensities in siTRIM8 samples versus the means quantified in the negative controls. The FCs range was fixed as |log2FC| ≥ 0.5 **(see Supplementary Methods for additional details)**.

### 4.5. Polysome profiling followed by RNA-sequencing

Media was aspirated from RPE cells (two 150 mm plates per condition). Plates were immediately placed on ice and washed with ice-cold 1x phosphate-buffered saline (PBS) containing cycloheximide (100 μg/mL, Sigma-Aldrich). Cells were lysed in 800 μL of polysome extraction buffer (10 mM Tris-HCl pH 7.4, 100 mM KCl, 10 mM MgCl2, 1% Triton X-100, 1 mM DTT, 10 U/mL RNaseOUT [Invitrogen], 100 μg/mL cycloheximide) and incubated on ice for 10 min. Lysates were cleared by centrifugation at 14,000 rpm at 4°C for 10 min. The supernatant was collected, and absorbance was measured at 260 nm using a NanoDrop. Between eight and ten optical density units were loaded onto a 10%–60% sucrose gradient formed by layering 6 mL of 10% sucrose over 6 mL of 60% sucrose prepared in polysome extraction buffer without Triton X-100 and containing 0.5 mM DTT, in a 12-mL tube (Polyallomer; Beckman Coulter). Gradients were prepared using a gradient maker (Gradient Master; Biocomp). Polysomes were separated by centrifugation at 37,000 rpm for 2 hours using a Beckman SW41 rotor. Twelve fractions of 920 μL were collected, and polysomes were monitored by measuring absorbance at 254 nm. RNA extraction was performed by adding TRI reagent (Merck T9424) at a 1:1 v/v ratio to the fractions, followed by the manufacturer’s protocol. Total RNA was processed using the SureSelect XT HS2 mRNA Library Preparation System kit (Agilent Technologies) according to the manufacturer’s instructions. The resulting cDNA libraries were quantified using the Qubit dsDNA HS Assay (Thermo Fisher) and analyzed by electrophoretic run on a TapeStation chip (Agilent Technologies). Sequencing was carried out on an Illumina NextSeq550 platform with a 2×75 paired-end run. For RNA-seq data analysis, alignment against the GRCh38 human genome was performed using STAR 2.5.4b^65^, with reads mapping to features quantified using HTSeq-count^66^. Differential expression analysis to identify differentially translated genes was conducted using DESeq2^67^ with R studio version 4.1.0.

### 4.6. Library preparation for BD Rhapsody scRNA-seq and sequencing with Illumina NextSeq 500

Sample preparation for mRNA Whole Transcriptome Analysis (WTA) followed the guidelines of the BD Single-Cell Multiplexing Kit (Cat. No. 633781). To distinguish between two conditions in the pool of cells, Sample Tag 05 was utilized for siRNA negative control RPE cells, while Sample Tag 06 was employed for TRIM8-silenced RPE cells. Transcriptomic analysis at the single-cell level was conducted using the BD Rhapsody Single-Cell Analysis System by BD Biosciences. For each sample, cell isolation was performed using the BD Single-Cell Multiplexing Kit (Cat. No. 633781, BD Biosciences), following the manufacturer’s guidelines. Cell viability and concentration were assessed using the BD Rhapsody Scanner system post-staining with Calcein AM (dilution ratio of 1:200; catalog number C1430, ThermoFisher) and DRAQ7™ (dilution ratio of 1:200; Cat. No. 564904, BD Biosciences), after a 5-minute incubation at 37°C. Cell counting was conducted using the Improved Neubauer Hemocytometer (INCYTO). Subsequently, cells from their respective samples were combined in a total volume of 650ml of cold BD Sample Buffer and loaded onto one BD Rhapsody cartridge with 10,000 pooled cells for the separation of individual cells. Isolation of single cells and synthesis of cDNA were achieved using the BD Rhapsody Express Single-Cell Analysis System, following the recommendations provided by BD Biosciences. The final suspension volume was determined based on the viable cells counted and captured, aiming to subsample and sequence approximately 4000 cells. Amplification of the whole transcriptome and Sample Tags was conducted using the BD Rhapsody Whole Transcriptome Amplification Kit (Cat. No. 633781, BD Biosciences), following the manufacturer’s protocol. Removal of undesired PCR products and small molecules was carried out with a cleanup step using AMPure XP Beckman magnetic beads (Cat. No. A63880, Beckman Coulter). The quantity and quality of DNA were assessed using the Qubit™ dsDNA HS Assay Kit (Cat. No. Q32851, ThermoFisher Scientific) and the Agilent 2200 TapeStation system, with the respective cartridge (Cat. No. 5067-5584). Two pools, RPE POOL1 (4000 cells) and RPE POOL2 (4000 cells), were generated. Each POOL comprised a mixed sample of *TRIM8*-silenced and siRNA control cells for their respective cell lines, distinguishable by sample tags as specified in the BD Rhapsody Library Preparation. To ensure rigor, each pool was run separately on the Illumina NextSeq 500 (NextSeq 500/550 High Output Kit v2.5; paired-end 2×75 bp sequencing). In each of the two runs, 4000 cells were targeted to be achieved, with an attempt to sequence a total of 8000 cells for further analysis. Sequencing was performed on the in-house NextSeq®500 System **(see Supplementary Methods for additional details)**.

### 4.7. Bioinformatic analysis of BD Rhapsody single-cell RNA sequencing (scRNA-seq) data

Raw single-cell RNA sequencing (scRNA-seq) data from the Illumina NextSeq 500 underwent initial quality assessment using Illumina Sequencing Analysis Viewer. Subsequently, FASTQ files underwent quality control analysis using FastQC. For single-cell analysis, FASTQ files were processed using the standard BD Rhapsody Pipeline, publicly available on the SevenBridges platform (BD Rhapsody™ WTA Analysis Pipeline). A data table containing molecule counts per gene per cell, based on Resampling-based Sequential Ensemble Clustering (RSEC) error correction, was generated. Principal Component Analysis (PCA) was employed to identify highly dispersed genes indicative of variance, capable of segregating biologically relevant populations within the data. PCA delineated two distinct clusters for control and *TRIM8*-silenced conditions, using sample tags assigned during library preparation (Illumina Sample Tag 05 for siRNA negative control RPE cells and Sample Tag 06 for TRIM8-silenced RPE cells). Additionally, t-SNE plots were utilized to preserve local structure in the high-dimensional dataset. PhenoGraph, a robust graph-based method^28^, was applied to t-SNE to identify subpopulations in high-dimensional single-cell data using 95 cell cycle candidate marker genes from Tirosh et al. (2016)^27^. Subsequent gene expression pattern analysis of cell cycle candidate marker genes among the clusters was conducted with the iCellR algorithm to identify unique clusters specific to G0/G1, S, and G2/M phases, based on the gene expression patterns of 95 cell cycle marker genes. Each of the three cellular clusters was further subdivided into *TRIM8*-silenced and control RPE cells based on the Illumina sample tags used during library preparation. Within each cluster, differentially expressed (DE) genes were subsequently identified between the *TRIM8*-silenced and control RPE populations of cells using the SeqGeq™ 1.7.0 bioinformatics analysis platform, tailored for BD Rhapsody scRNA-seq data analysis.

### 4.8. Cell synchronization, BrdU/7-AAD cell cycle assay, and flow cytometry

RPE cells were synchronized in mitosis using a palbociclib–nocodazole block method and subsequently harvested at the G1 phase after 75 minutes of release from the palbociclib–nocodazole block, as described by Scott et al. (2020)^68^. After synchronization, RPE cells were seeded, and TRIM8-EGFP overexpression was induced six hours post-seeding using the Lipofectamine™ LTX-PLUS protocol, following the manufacturer’s instructions, at a concentration of 5 μM. Following a two-hour interval post-seeding, ensuring a total time interval of 8 hours from seeding, 10 µM BrdU was added to the cells. The duration of cell cycle phases in RPE cells has been well-standardized by Chao et al. (2019)^69^, with durations of ∼7.9 hours for G1, ∼7.6 hours for S, ∼3.4 hours for G2, and ∼0.5 hours for M-phase. In accordance with these established time points, BrdU-treated TRIM8-EGFP-positive RPE cells and BrdU-treated RPE cells lacking TRIM8-EGFP were simultaneously collected at the conclusion of the S-phase to maximize the likelihood of capturing BrdU-labeled cells. BrdU serves as a synthetic nucleoside analogue of thymidine and is indicative of newly synthesized DNA during the S-phase^70^. The collected cells were then subjected to the BrdU/7-AAD staining protocol following the manufacturer’s instructions provided by the APC BrdU Flow Kit (Cat. No. 557892, BD Biosciences). Subsequently, cells were sorted using flow cytometer based on GFP detection, allowing differentiation between untransfected and transfected cells. Viable cells were distinguished by negative staining for DAPI (Cat. No. D9542, Sigma-Aldrich), followed by debris exclusion and singlet gating based on forward and side scatter. The different cell cycle phases were determined by analysing the correlated expression of total DNA and incorporated BrdU levels, along with fluorescence signals from 7-AAD. Flow cytometry was performed on BD FACSCanto™ II and flow cytometry data was analysed using FlowJo™ software (BD Biosciences).

### 4.9. Confocal microscopy and immunofluorescence study

For the localisation of TRIM8 at primary cilium, RPE cells were subjected to serum starvation for 48 hours in DMEM medium devoid of fetal bovine serum (FBS). The primary cilium was identified using an anti-acetylated alpha tubulin (acetyl K40) antibody (ab125356), which served as the marker for primary cilium visualization. For the TRIM8-CEP170 colocalization study, HeLa cells were synchronized with a 12-hour double block using thymidine at a concentration of 2mM. For the immunofluorescence study, cells were fixed in ice-cold methanol for 15 minutes at +4°C. Blocking was performed in PBS containing 1% BSA for 1 hour at room temperature (RT), followed by incubation with primary antibodies (1° Ab) in PBS supplemented with 0.1% BSA for 2 hours at RT. Primary antibodies used in the experiment included rabbit anti-CEP170 N-term at a dilution of 1:600 (sourced from Guarguaglini et al, 2005), anti-TRIM8 at a dilution of 1:200 (sc-398878, Santa Cruz), rabbit anti-TRIM8 at a dilution of 1:100 (HPA023561, Sigma), and mouse anti-Y-tubulin at a dilution of 1:1000 (T-6557, Sigma). Following primary antibody incubation, cells were incubated with secondary antibodies (2° Ab) in PBS supplemented with 0.1% BSA for 1 hour at RT. Additionally, cells were stained with 200 nM DAPI for 5 minutes at RT to visualize nuclei. After completion of the staining protocol, cells were imaged using Zeiss LSM700 Confocal Microscope to examine the localization of the target proteins.

## ACKNOWLEDGEMENT

The authors thank Giulia Guarguaglini (Sapienza University of Rome, Italy) for providing hTERT RPE-1 cells and the CEP170 antibody for the co-localization study.

## SUPPLEMENTARY DATA

Supplementary data will be available in the peer-reviewed publication or upon request to the corresponding author.

